# Bacterial Lytic Polysaccharide Monooxygenases negatively affect insect larvae growth

**DOI:** 10.1101/2024.12.10.627774

**Authors:** Jakub Baranek, Filip Wojtkowiak, Kinga Belińska, Paulina Wojtkowiak, Andrzej Zielezinski

## Abstract

**BACKGROUND:** Lytic Polysaccharide Monooxygenases (LPMOs) are recently discovered redox enzymes broadly distributed across various taxa, including bacteria, fungi, and viruses. LPMO proteins can oxidatively cleave the glycosidic chains found in carbohydrate polymers, allowing efficient decomposition of recalcitrant polysaccharides such as chitin or cellulose. While LPMOs have primarily been studied for their role in biomass conversion, other biological roles of these enzymes are not well understood.

**RESULTS:** Here, we assess the influence of four different LMPOs, derived from *Bacillus thuringiensis* and *Serratia marcescens* bacteria, on *Spodoptera exigua* larvae. The tested proteins administered *per os* resulted in significant growth impairment, with the mean body weight of the treated larvae being lower than the control group by 30-69 percentage points, depending on the specific LPMO.

**CONCLUSION:** This work indicates that LPMO enzymes are capable of insect larvae development inhibition and thus may play an important role in insect pathogenesis caused by bacteria. Moreover, the results obtained in this study are an early sign of the potential use of LPMOs in future pest management strategies.

## Introduction

Lytic Polysaccharide Monooxygenases (LPMOs) are copper-dependent redox enzymes capable of oxidative cleavage of glycosidic bonds in recalcitrant carbohydrates such as chitin or cellulose. LPMOs along with various enzymes such as cellulases, chitinases and other glycoside hydrolases play a key role in the depolymerization of polysaccharides and therefore are crucial in biomass degradation and the carbon cycle on earth. The biological importance of LPMOs is reflected in a great abundance of genes encoding these proteins, which are found in various species, representing viruses, bacteria, and fungi ^1^.

Since the discovery of LPMOs in the early 21^st^ century, numerous studies have been performed to characterize LPMOs, and these findings have been comprehensively reviewed in recent works ^1–4^. Proteins currently known as LPMOs are classified in the Carbohydrate-Active enZYme (CAZy) database ^5^ as auxiliary activity (AA) families AA9, AA10, AA11, AA13, AA14, and AA15. However, there is a wide range of diversity among and within the families of LPMOs in terms of their preference for different substrates (e.g., cellulose, chitin, starch, xyloglucan), as well as their sequence length, structural features, taxonomic association, and kinetics ^1,3^. Despite the growing knowledge regarding LPMOs, many aspects remain uncertain, including their exact mode of action and biological roles, which apart from biomass degradation may also involve pathogenesis, body development, or food digestion ^1^.

It has been shown that some of LPMOs oxidize crystalline forms of chitin ^6,7^, and they act synergistically with chitinases – classical glycoside hydrolases, capable of degradation of chitin in its soluble form. As a result, mixtures of glycoside hydrolases and LPMOs achieve higher polysaccharide degradation activity than any of the components alone ^8–10^. Many invertebrates, such as insects, use chitin as the main structural component of their exoskeleton and as the lining of their intestine (peritrophic matrix), which plays a crucial role in digestion and protecting the insect gut from pathogens and harmful molecules like toxins ^11^. Various chitinases were proven to contribute to the microbial pathogenicity towards insects by disrupting the peritrophic matrix. Due to their insecticidal properties, chitinases are used to enhance the protection of crops against pests, thus serving as biocontrol agents ^12^. It is probable that certain LPMOs may also exert adverse effects on invertebrates (e.g., by peritrophic matrix function impairment) and could potentially be used in the future as next-generation bioinsecticides, either independently or in combination with other bioactive molecules.

However, the direct influence of LPMOs on invertebrates and their possible implementation as bioinsecticides has been tested only marginally. First, it was that demonstrated very weak synergistic interaction occur between *Bacillus thuringiensis* 21 kDa LPMO designated Bt-CBP21 and *Bacillus thuringiensis* Cry1Ac pesticidal protein, in *Helicoverpa armigera* larvae ^13^. Next, the upregulation of the gene encoding LPMO in *Lysinibacillus sphaericus* during infection of *Aedes aegypti* has been shown ^14^. Finally, in a recent study it was found that the LPMO-deficient mutant of *B. thuringiensis* HD73 had 10-fold lower insecticidal activity against *Ostrinia furnacalis* compared to the wild-type strain, and LPMO improved the adherence of *B. thuringiensis* HD73 vegetative cells to the peritrophic membrane of the insect ^15^. Although the above findings suggest LPMOs may be the active components of the bacterial entomopathogenic arsenal, many questions remain to be answered, including whether LPMOs act only as accessory components (synergizing other insecticidal molecules or improving the adherence of vegetative cells of entomopathogens) or if they can exert negative effects on invertebrate organisms alone. Moreover, the previous studies were limited to a few closely related species that belong to the *Bacillaceae* family within the Firmicutes phylum, and thus it remains unknown if LPMOs produced by other microorganisms (i. e., gram-negative bacteria) will also have negative effects on invertebrates.

In this study, we examine the effects of various LPMO enzymes when administered individually *per os*, on larvae of the beet armyworm (*Spodoptera exigua* Hübn; Lepidoptera: Noctuidae), an economically important and polyphagous pest species. The tested LPMOs included four heterologously-expressed proteins, two proteins from gram-positive entomopathogenic bacterium *Bacillus thuringiensis* and the other two from the gram-negative microorganism *Serratia marcescens*, which is known for its efficient chitinolytic properties. The results obtained in this work provide new insights into the role of LPMOs in insecticidal action and hints at the potential use of LPMOs in biocontrol strategies.

## Materials and Methods

### In silico analysis

The LPMO proteins used for sequence comparison were selected based on their proven chitin-targeting properties reported in source articles ^8,10,16–27^. Functional domains of LPMO proteins analyzed in this study were detected using InterPro ^28^ and dbCAN2 ^29^ databases. The amino acid sequences of LPMOs were aligned using needle (with default parameters) from the EMBOSS package v. 6.6.0.0 ^30^.

### Protein expression

Genes encoding BtLPMO10_50, BtLPMO10_24, and 74 kDa chitinase from *B. thuringiensis* (strain MPU B7) and genes encoding SmLPMO10_21 and SmLPMO10_52 from *S. marcescens* (strain Sm5), were PCR amplified and subsequently cloned into linearized pLATE11 expression vectors. Heterologous expression was carried out in *E. coli* BL21 (DE3) codon+ cells. The proteins were extracted from cell lysates, quantified and frozen at -75°C until further use. The detailed procedure of obtaining proteins is described in **Supplementary file 1**. Simultaneously, the *B. thuringiensis* Cry2Ab pesticidal protein was produced, as reported previously ^31^, serving as a positive control in this study.

### Insect bioassays

Diet surface contamination bioassays were performed on the first instar (2-3-day-old) *S. exigua* larvae using multi-well plates containing a lepidopteran-dedicated diet ^32^. Diet in each well was covered with a given LPMO protein or chitinase at a concentration of 3000 ng/cm^2^.

For each tested protein, 22 lepidopteran larvae were assigned, and all bioassays were done in three independent repetitions (in total, 66 larvae per assayed protein). Cleared lysate from non-transformed *E. coli* BL21 (DE3) codon plus cells was used as a negative control, whereas *B. thuringiensis* Cry2Ab protoxin at a concentration of 300 ng/cm^2^ was used as a positive control.

To determine larval growth inhibition caused by tested proteins, both the larvae surviving LPMO treatments and the control insects were weighed using 0.1 mg precision, seven days after treatment. The mean body weight of the treated larvae was expressed as a percent of negative control larvae body weight. The mortality was also scored seven days after treatment, and the insect was considered dead when no movement was seen after slight agitation with a brush. Because the mean mortality in the controls was only ∼5%, the correction of the mortality in the treated larvae was omitted. In both approaches, significant differences between the treatment larvae and the control were analyzed in R v. 4.2.2 by one-way analysis of variance (ANOVA), followed by Tukey’s test, with a significance level of *P* < 0.05.

## Results

### LPMO sequence analysis

The four LPMO proteins introduced in this study belong to the auxiliary activity 10 family (AA10) of the carbohydrate-active enzymes, similarly to previously studied chitin-targeting LPMO enzymes (Fig. 1). Although all four LPMOs share a single AA10 protein domain, they represent three different domain architectures, suggesting their functional diversity.

**Fig. 1.**
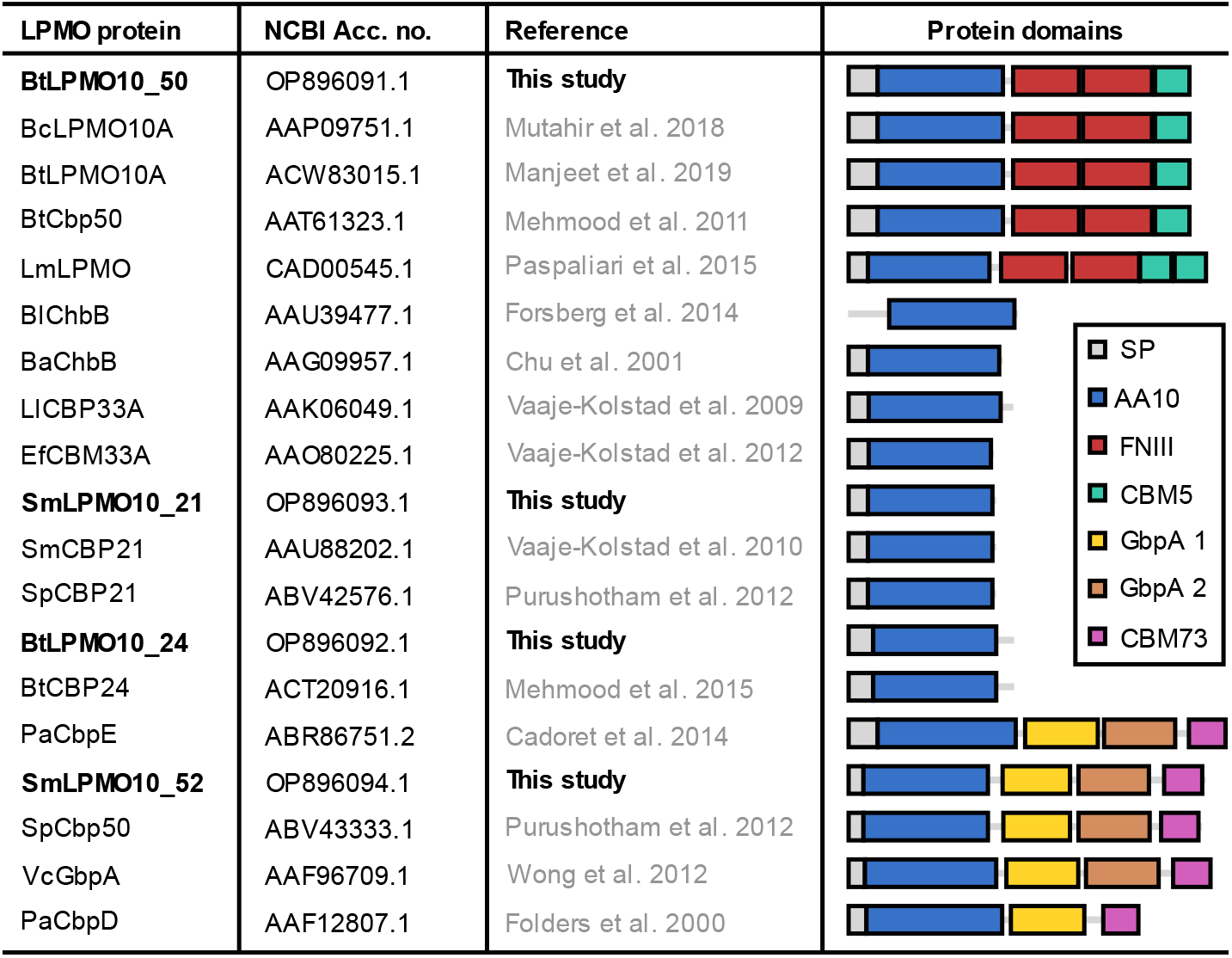
Comparison of four LPMO proteins characterized in this study to other proteins with known chitin-targeting LPMOs. The protein domains and functional motifs are denoted as: SP – signal peptide; AA10 – auxiliary activity family 10 (AA10) domain; FNIII – fibronectin type III; CBM5 –carbohydrate binding module family 5; GbpA 1 - N-Acetylglucosamine binding protein A type 1; GbpA 2 – N-Acetylglucosamine binding protein A type 2; CBM73 - carbohydrate binding module family 73. Abbreviations of bacterial species used in LPMO names are: Ba – *Bacillus amyloliquefaciens*; Bc – *B. cereus*; Bl – *B. licheniformis*; Bt – *B. thuringiensis;* Ef – *Enterococcus faecalis*; Ll – *Lactococcus lactis*; Lm - *Listeria monocytogenes;* Pa – *Pseudomonas aeruginosa*; Sm – *Serratia marcescens*; Sp – *S. proteamaculans*; Vc – *Vibrio cholerae*

Specifically, BtLPMO10_24 and SmLPMO10_21 enzymes feature only AA10 domain. The BtLPMO10_50 contains the AA10 domain, two fibronectin type III domains, and carbohydrate-binding module type 5. Finally, SmLPMO10_52 is comprised of the AA10 domain, two N-Acetylglucosamine binding protein A domains, and carbohydrate-binding module type 73.

The amino acid sequences of four enzymes studied in this work show high resemblance to other LPMOs with well-documented enzymatic and functional properties (**Supplementary file 2**). For example, BtLPMO10_50 has 98.2% sequence identity with BcLPMO10A from *Bacillus cereus* ATCC 14579, which has proven enzymatic activity on various forms of chitin and synergizes action of different chitinases ^10^. BtLPMO10_24 has 93.2% sequence identity with *B. thuringiensis* serovar *konkukian* BtCBP24, which has been shown to bind to chitin, act synergistically with chitinases characteristic for *B. thuringiensis* and *Serratia proteamaculans*, and display significant antifungal properties ^8^. SmLPMO10_21 has 99.5% sequence identity with *S. marcescens* SmCBP21, which is capable of oxidative cleavage of chitin chains and boosting various chitinolytic enzymes produced by this bacterium ^22,25,33^.

Only in the case of SmLPMO10_52, the similarity to LPMOs with documented chitin activity is much lower. Examples of the most similar LPMOs to SmLPMO10_52 include: (i) SpCbp50 derived from *S. proteamaculans* (57% sequence identity), which acts synergistically with various chitinolytic enzymes produced by this microbe ^23^; and (ii) VcGbpA from *Vibrio cholerae* (41.8% sequence identity), an enigmatic LPMO sharing properties associated with both chitin degradation ^22,27,34^ as well as pathogenicity ^35–38^.

### Insect bioassays

To determine the biological activity of individual *B. thuringiensis* and *S. marcescens* LPMOs towards insects, the four enzymes, featuring various domain architectures were administered to first instar larvae of *S. exigua* using a surface contamination assay. For comparison, the *B. thuringiensis* 74 kDa chitinase (GenBank Acc. No. OP948078.1) was also included in the test due to its reported insecticidal and nematicidal properties ^39,40^. Additionally, the *B. thuringiensis* Cry2Ab insecticidal protein was included in the bioassays as a positive control, as it has been shown to exert negative effects against various lepidopteran larvae, including *S. exigua* ^31,41,42^.

The results show that the larvae treated with each of the tested LPMOs had a mean body weight lower by 30-69 percentage points than in control (Fig. 2a). These differences were statistically significant for two LPMOs, derived from *S. marcescens* (SmLPMO10_21 and SmLPMO10_52). Moreover, larvae fed with diet containing the two latter enzymes had lower mean body weight by 31-41 percentage points compared to larvae fed with BtChi_74 chitinase. During this part of the study, we also monitored insect mortality and observed that three of the tested LPMOs (namely BtLPMO10_50, BtLPMO10_24, and SmLPMO21) increased the mean mortality rate by 4-11 percentage points compared to the control group, and 3-10 percentage points compared to the chitinase-treated larvae, although the differences were not statistically significant (Fig. 2b). In summary, the bacterial LPMOs, administered *per os*, cause adverse effects in *S. exigua* larvae development, and these effects are comparable or higher (depending on LPMO) than those caused by a bacterial chitinase, which has previously been shown to have biopesticidal properties.

**Fig. 2.**
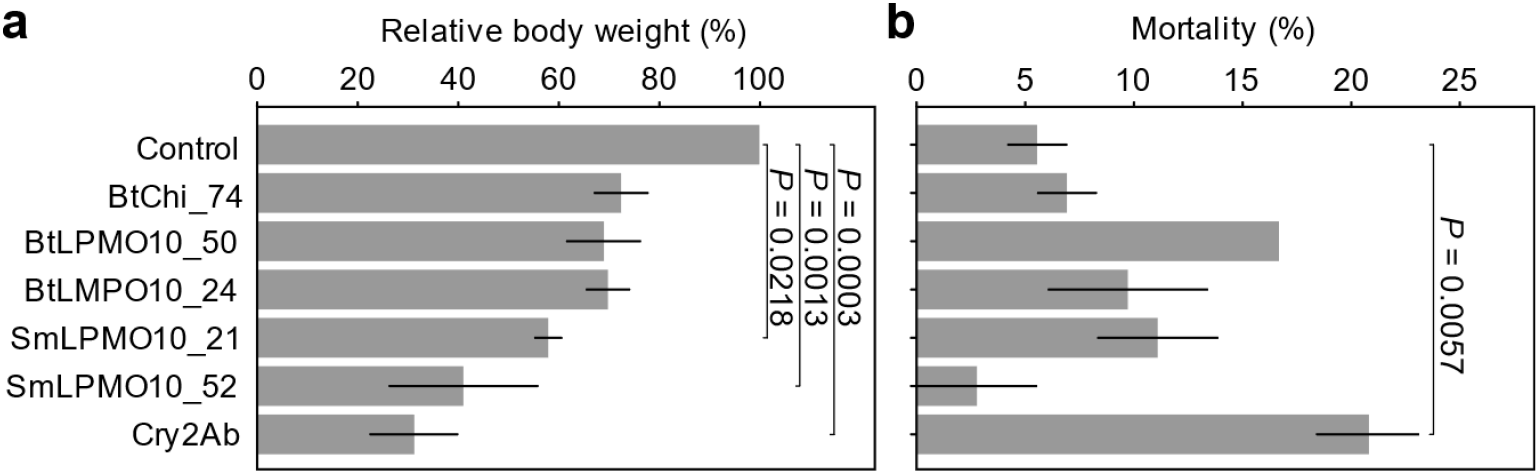
Relative body weight (a) and mortality (b) of *Spodoptera exigua* first instar larvae after treatment with *Bacillus thuringiensis* (Bt) and *Serratia marcescens* (Sm) LPMO proteins, administered *per os* at a concentration of 3000 ng/cm^2^. The results were scored seven days after treatment. For comparison, the effects of *B. thuringiensis* 74 kDa chitinase (BtChi_74) and Cry2Ab insecticidal protein on the larvae have also been included. The *P*-values are shown for statistically significant differences (*P* < 0.05) between the treated larvae and negative control. Error bars indicate standard error of the mean

## Discussion

This study shows that LPMO enzymes can disrupt larvae development when administered to the test insect independently. This is an essential addition to the limited knowledge regarding LPMOs because the results of previous studies suggested that these enzymes can act only by synergizing the insecticidal activity of *B. thuringiensis or L. sphaericus* ^13–15^. For example, the work performed by Qin et al. (2020), shows the role of LPMO protein (designated initially as CBPA; GenBank Acc. No. YP_007422288.1) in increasing *B. thuringiensis* insecticidal activity by improvement of the bacterial adherence to insect peritrophic membrane. Our results suggest that at least one different mode of action exists regarding LPMO enzymes, not involving any living bacterial cells. We can speculate that the LPMO proteins may negatively affect a specific structure (or structures) of the insect body when individually administered *per os*. The most probable target is the peritrophic membrane, which may be enzymatically disrupted by LPMOs, resulting in improper function of the gut and digestive processes, and finally in growth impairment.

Noteworthy is the fact that *S. exigua* larvae growth inhibition has been caused by two LPMOs derived from *S. marcescens*. This is the first report on negative effects caused by LPMOs characteristic for gram-negative bacterium, towards insects. Although *S. marcescens* is mainly considered a pathogen of animals ^43,44^ and humans ^45^, it can also infect plants ^46,47^ and some strains of this species have been shown to infect insects ^48–50^. This work shows that *S. marcescens* LPMOs negatively affect insect larvae and thus may be involved in insect infections caused by this microbe. The influence of LPMO proteins on *S. marcescens* pathogenicity to other hosts also cannot be excluded.

To date, numerous studies have been performed, showing the detrimental effects of various bacterial chitinases on different invertebrate species ^12^. The most probable explanation of the chitinase biocidal effects is the disruption of the peritrophic matrix, which protects insects or nematodes from foodborne pathogens and toxins ^11^. High implementation potential has been noticed in chitinases, which may be used in pest biocontrol strategies ^12^. Usually, these enzymes do not cause insect mortality alone but have been shown as synergistic molecules, which have the ability to increase the pesticidal activity of *B. thuringiensis* Cry proteins ^39,40,51–53^. Such is the case of 74 kDa chitinase, derived from *B. thuringiensis*, which has been proven to boost insecticidal and nematicidal properties of *B. thuringiensis* ^39,40^. The results of our study demonstrate that two *S. marcescens* LPMOs exert a higher growth-inhibition effect on *S. exigua* larvae than the 74 kDa *B. thuringiensis* chitinase. Therefore, it is possible that LPMO proteins may have higher implementation potential in biocontrol strategies than the classical chitinolytic enzymes. Nonetheless, it would require a much broader approach to perform a thorough comparison of these two groups, for example by implementing bioassays, which involve different chitinases tested along with LPMOs at a wide range of concentrations. Moreover, to better estimate the potential of LPMOs in biocontrol strategies, these enzymes should be assessed for their synergistic interactions with chitinases and other biopesticidal agents (such *B. thuringiensis* Cry and Vip toxins) against decent number of distinct pest species.

In conclusion, bacterial LPMO enzymes cause insect larvae growth impairment, when administered *per os*, without the presence of additional factors such as bacterial cells or insecticidal toxins. This indicates that LPMOs may play a role (at least auxiliary) in pathogenicity of bacteria towards insects. Furthermore, this work is an early prognostic of the possible use of LPMOs as next-generation pest killers used in biological pest management, including Integrated Pest Management.

## Acknowledgements

This work was supported by the National Science Centre, Poland [grant no. 2018/02/X/NZ9/00232] to J.B. The authors would like to thank Halina Paetz, Anna Sufryd-Sikora and Renata Sikorska for their technical support. We are also grateful to Joanna Nowicka-Baranek for the linguistic improvements to the manuscript.

## Declarations

### Ethics approval and consent to participate

This article does not contain any studies with human participants or animals (vertebrates) performed by any of the authors.

### Consent for publication

Not applicable.

### Availability of data and materials

The data that support the findings of this study are available from the corresponding author upon reasonable request.

## References

1 Forsberg Z, Sørlie M, Petrović D, Courtade G, Aachmann FL, Vaaje-Kolstad G, et al., Polysaccharide degradation by lytic polysaccharide monooxygenases, Curr Opin Struct Biol 59:54–64 (2019).

2 Johansen KS, Lytic polysaccharide monooxygenases: the microbial power tool for lignocellulose degradation, Trends Plant Sci 21:926–936, Elsevier Ltd (2016).

3 Courtade G and Aachmann FL, Chitin-active lytic polysaccharide monooxygenases, ed. by Yang Q and Fukamizo T, Targeting chitin-containing organisms, Springer, Singapore, pp. 115–129 (2019).

4 Eijsink VGH, Petrovic D, Forsberg Z, Mekasha S, Røhr ÅK, Várnai A, et al., On the functional characterization of lytic polysaccharide monooxygenases (LPMOs), Biotechnol Biofuels 12:1–16, BioMed Central (2019).

5 Drula E, Garron ML, Dogan S, Lombard V, Henrissat B, and Terrapon N, The carbohydrate-active enzyme database: functions and literature, Nucleic Acids Res 50:D571–D577, Oxford University Press (2022).

6 Bissaro B, Isaksen I, Vaaje-Kolstad G, Eijsink VGH, and Røhr ÅK, How a lytic polysaccharide monooxygenase binds crystalline chitin, Biochemistry 57:1893–1906 (2018).

7 Kuusk S, Kont R, Kuusk P, Heering A, Sørlie M, Bissaro B, et al., Kinetic insights into the role of the reductant in H2O2-driven degradation of chitin by a bacterial lytic polysaccharide monooxygenase, Journal of Biological Chemistry 294:1516–1528 (2019).

8 Mehmood MA, Latif M, Hussain K, Gull M, Latif F, and Rajoka MI, Heterologous expression of the antifungal-chitin binding protein CBP24 from Bacillus thuringiensis and its synergistic action with bacterial chitinases, Protein Pept Lett 22:39–44 (2015).

9 Mehmood MA, Hussain K, Latif F, Tabassum MR, Gull M, Gill SS, et al., Synergistic action of the antifungal β-chitin binding protein CBP50 from Bacillus thuringiensis with bacterial chitinases, Curr Proteomics 11:23–26, Bentham Science Publishers B.V. (2014).

10 Mutahir Z, Mekasha S, Loose JSM, Abbas F, Vaaje-Kolstad G, Eijsink VGH, et al., Characterization and synergistic action of a tetra-modular lytic polysaccharide monooxygenase from Bacillus cereus, FEBS Lett 592:2562–2571 (2018).

11 Hegedus DD, Toprak U, and Erlandson M, Peritrophic matrix formation, J Insect Physiol 117:103898, Elsevier (2019).

12 Berini F, Katz C, Gruzdev N, Casartelli M, Tettamanti G, and Marinelli F, Microbial and viral chitinases: attractive biopesticides for integrated pest management, Biotechnol Adv 36:818–838, Elsevier (2018).

13 Arora N, Sachdev B, Gupta R, Vimala Y, and Bhatnagar RK, Characterization of a chitin-binding protein from Bacillus thuringiensis HD-1, PLoS One 8:66603, International Center for Genetic Engineering and Biotechnology, Aruna Asaf Ali Marg, New Delhi, India (2013).

14 Rojas-Pinzón PA and Dussán J, Contribution of Lysinibacillus sphaericus hemolysin and chitin-binding protein in entomopathogenic activity against insecticide resistant Aedes aegypti, World J Microbiol Biotechnol 33:1–9, Springer Netherlands (2017).

15 Qin J, Tong Z, Zhan Y, Buisson C, Song F, He K, et al., A Bacillus thuringiensis chitin-binding protein is involved in insect peritrophic matrix adhesion and takes part in the infection process, Toxins (Basel) 12 (2020).

16 Cadoret F, Ball G, Douzi B, and Voulhoux R, Txc, a new type II secretion system of Pseudomonas aeruginosa strain PA7, is regulated by the TtsS/TtsR two-component system and directs specific secretion of the CbpE chitin-binding protein, J Bacteriol 196:2376–2386 (2014).

17 Chu HH, Hoang V, Hofemeister J, and Schrempf H, A Bacillus amyloliquefaciens ChbB protein binds β-and α-chitin and has homologues in related strains, Microbiology (N Y) 147:1793–1803 (2001).

18 Folders J, Tommassen J, van Loon LC, and Bitter W, Identification of a chitin-binding protein secreted by Pseudomonas aeruginosa, J Bacteriol 182:1257–1263 (2000).

19 Forsberg Z, Røhr ÅK, Mekasha S, Andersson KK, Eijsink VGH, Vaaje-Kolstad G, et al., Comparative study of two chitin-active and two cellulose-active AA10-type lytic polysaccharide monooxygenases, Biochemistry 53:1647–1656 (2014).

20 Manjeet K, Madhuprakash J, Mormann M, Moerschbacher BM, and Podile AR, A carbohydrate binding module-5 is essential for oxidative cleavage of chitin by a multi-modular lytic polysaccharide monooxygenase from Bacillus thuringiensis serovar kurstaki, Int J Biol Macromol 127:649–656, Elsevier B.V. (2019).

21 Mehmood MA, Xiao X, Hafeez FY, Gai Y, and Wang F, Molecular characterization of the modular chitin binding protein Cbp50 from Bacillus thuringiensis serovar konkukian, Antonie Van Leeuwenhoek 100:445–453 (2011).

22 Paspaliari DK, Loose JSM, Larsen MH, and Vaaje-Kolstad G, Listeria monocytogenes has a functional chitinolytic system and an active lytic polysaccharide monooxygenase, FEBS Journal 282:921–936 (2015).

23 Purushotham P, Arun PVPS, Prakash JSS, and Podile AR, Chitin binding proteins act synergistically with chitinases in Serratia proteamaculans 568, PLoS One 7:1–10 (2012).

24 Vaaje-Kolstad G, Bunæs AC, Mathiesen G, and Eijsink VGH, The chitinolytic system of Lactococcus lactis ssp. lactis comprises a nonprocessive chitinase and a chitin-binding protein that promotes the degradation of α- And β-chitin, FEBS Journal 276:2402–2415 (2009).

25 Vaaje-Kolstad G, Westereng B, Horn SJ, Liu Z, Zhai H, Sørlie M, et al., An Oxidative Enzyme Boosting the Enzymatic Conversion of Recalcitrant Polysaccharides, Science (1979) 330:219–222 (2010).

26 Vaaje-Kolstad G, Bøhle LA, Gåseidnes S, Dalhus B, Bjørås M, Mathiesen G, et al., Characterization of the chitinolytic machinery of Enterococcus faecalis V583 and high-resolution structure of its oxidative CBM33 enzyme, J Mol Biol 416:239–254, Elsevier Ltd (2012).

27 Wong E, Vaaje-Kolstad G, Ghosh A, Hurtado-Guerrero R, Konarev P v., Ibrahim AFM, et al., The Vibrio cholerae colonization factor GbpA possesses a modular structure that governs binding to different host surfaces, PLoS Pathog 8:1–12 (2012).

28 Paysan-Lafosse T, Blum M, Chuguransky S, Grego T, Pinto BL, Salazar GA, et al., InterPro in 2022, Nucleic Acids Res:1–10 (2022).

29 Zhang H, Yohe T, Huang L, Entwistle S, Wu P, Yang Z, et al., DbCAN2: a meta server for automated carbohydrate-active enzyme annotation, Nucleic Acids Res 46:W95–W101, Oxford University Press (2018).

30 Rice P, Longden L, and Bleasby A, EMBOSS: the european molecular biology open software suite, Trends in Genetics 16:276–277 (2000).

31 Baranek J, Banaszak M, Kaznowski A, and Lorent D, A novel Bacillus thuringiensis Cry9Ea‐like protein with high insecticidal activity towards Cydia pomonella larvae, Pest Manag Sci 77:1401–1408 (2021).

32 McGuire MR, Galan-Wong LJ, and Tamez-Guerra P, Bioassay of Bacillus thuringiensis against lepidopteran larvae, ed. by Lacey LA, Manual of Techniques in Insect Pathology, Academic Press, San Diego, California, USA, pp. 91–99 (1997).

33 Nakagawa YS, Eijsink VGH, Totani K, and Vaaje-Kolstad G, Conversion of α-chitin substrates with varying particle size and crystallinity reveals substrate preferences of the chitinases and lytic polysaccharide monooxygenase of Serratia marcescens, J Agric Food Chem 61:11061–11066 (2013).

34 Loose JSM, Forsberg Z, Fraaije MW, Eijsink VGH, and Vaaje-Kolstad G, A rapid quantitative activity assay shows that the Vibrio cholerae colonization factor GbpA is an active lytic polysaccharide monooxygenase, FEBS Lett 588:3435–3440 (2014).

35 Kirn TJ, Jude BA, and Taylor RK, A colonization factor links Vibrio cholerae environmental survival and human infection, Nature 438:863–866 (2005).

36 Zampini M, Pruzzo C, Bondre VP, Tarsi R, Cosmo M, Bacciaglia A, et al., Vibrio cholerae persistence in aquatic environments and colonization of intestinal cells: involvement of a common adhesion mechanism, FEMS Microbiol Lett 244:267–273 (2005).

37 Bhowmick R, Ghosal A, Das B, Koley H, Saha DR, Ganguly S, et al., Intestinal adherence of Vibrio cholerae involves a coordinated interaction between colonization factor GbpA and mucin, Infect Immun 76:4968–4977 (2008).

38 Jude BA, Martinez RM, Skorupski K, and Taylor RK, Levels of the secreted Vibrio cholerae attachment factor GbpA are modulated by quorum-sensing-induced proteolysis, J Bacteriol 191:6911–6917 (2009).

39 Ni H, Zeng S, Qin X, Sun X, Zhang S, Zhao X, et al., Molecular docking and site-directed mutagenesis of a Bacillus thuringiensis chitinase to improve chitinolytic, synergistic lepidopteran-larvicidal and nematicidal activities, Int J Biol Sci 11:304–315 (2015).

40 Qin X, Xiang X, Sun X, Ni H, and Li L, Preparation of nanoscale Bacillus thuringiensis chitinases using silica nanoparticles for nematicide delivery, Int J Biol Macromol 82:13–21, Elsevier B.V. (2016).

41 Lu Q, Cao GC, Zhang LL, Liang GM, Gao XW, Zhang YJ, et al., The binding characterization of Cry insecticidal proteins to the brush border membrane vesicles of Helicoverpa armigera, Spodoptera exigua, Spodoptera litura and Agrotis ipsilon, J Integr Agric 12:1598–1605, Chinese Academy of Agricultural Sciences (2013).

42 Tabashnik BE, Fabrick JA, Unnithan GC, Yelich AJ, Masson L, Zhang J, et al., Efficacy of genetically modified Bt toxins alone and in combinations against pink bollworm resistant to Cry1Ac and Cry2Ab, PLoS One 8:e80496, Department of Entomology, University of Arizona, Tucson, AZ, United States (2013).

43 Das AM, Paranjape VL, and Pitt TL, Serratia marcescens infection associated with early abortion in cows and buffaloes, Epidemiol Infect 101:143–149 (1988).

44 Saidenberg ABS, Teixeira RHF, Astolfi-Ferreira CS, Knöbl T, and Piantino Ferreira AJ, Serratia marcescens infection in a swallow-tailed hummingbird, J Wildl Dis 43:107–110 (2007).

45 Hejazi A and Falkiner FR, Serratia marcescens, J Med Microbiol 46:903–912 (1997).

46 Bruton BD, Mitchell F, Fletcher J, Pair SD, Wayadande A, Melcher U, et al., Serratia marcescens, a phloem-colonizing, squash bug-transmitted bacterium: causal agent of cucurbit yellow vine disease, Plant Dis 87:937–944 (2003).

47 Wang XQ, Bi T, Li XD, Zhang LQ, and Lu SE, First report of corn whorl rot caused by Serratia marcescens in China, Journal of Phytopathology 163:1059–1063 (2015).

48 Pineda-Castellanos ML, Rodríguez-Segura Z, Villalobos FJ, Hernández L, Lina L, and Nuñez-Valdez ME, Pathogenicity of isolates of Serratia marcescens towards larvae of the scarab Phyllophaga blanchardi (Coleoptera), Pathogens 4:210–228 (2015).

49 Fu R, Zhou L, Feng K, Lu X, Luo J, and Tang F, Effects of Serratia marcescens (SM1) and its interaction with common biocontrol agents on the termite, Odontotermes formosanus (Shiraki), J For Res (Harbin) 32:1263–1267, Springer Berlin Heidelberg (2021).

50 Zhang P, Zhao Q, Ma X, and Ma L, Pathogenicity of Serratia marcescens to hazelnut weevil (Curculio dieckmanni), J For Res (Harbin) 32:409–417, Springer Berlin Heidelberg (2021).

51 Regev A, Keller M, Strizhov N, Sneh B, Prudovsky E, Chet I, et al., Synergistic activity of a Bacillus thuringiensis delta-endotoxin and a bacterial endochitinase against Spodoptera littoralis larvae., Appl Environ Microbiol 62:3581–3586 (1996).

52 Hu SB, Zhang YM, Li WP, Xia LQ, Sun YJ, Liu P, et al., Efficient constitutive expression of chitinase in the mother cell of Bacillus thuringiensis and its potential to enhance the toxicity of Cry1Ac protoxin, Appl Microbiol Biotechnol 82:1157–1167, Berlin/Heidelberg : Springer-Verlag (2009).

53 González-Ponce KS, Casados-Vázquez LE, Lozano-Sotomayor P, Bideshi DK, del Rincón-Castro MC, and Barboza-Corona JE, Expression of ChiA74Δsp and its truncated versions in Bacillus thuringiensis HD1 using a vegetative promoter maintains the integrity and toxicity of native Cry1A toxins, Int J Biol Macromol 124:80–87, Elsevier B.V. (2019).

