## Supplementary file 1 for "Bacterial Lytic Polysaccharide Monooxygenases negatively affect insect larvae growth"

***Bacillus thuringiensis*** **and** ***Serratia marcescens*** **Lytic Polysaccharide Monooxygenases exert negative effects on** ***Spodoptera exigua*** **larvae**

Jakub Baranek[1*], Filip Wojtkowiak[1], Kinga Belińska[1], Paulina Wojtkowiak[1], Andrzej Zielezinski[2]

[1]Department of Microbiology, Faculty of Biology, Adam Mickiewicz University in Poznań, Uniwersytetu Poznańskiego 6, 61-614, Poznań, Poland

[2]Department of Computational Biology, Faculty of Biology, Adam Mickiewicz University in Poznań, Uniwersytetu Poznańskiego 6, 61-614, Poznań, Poland

### Materials and Methods

Cloning, expression and extraction of recombinant proteins

Genomic DNAs from the *B. thuringiensis* (strain MPU B7) and *S. marcescens* (strain Sm5) were isolated using GeneMATRIX Bacterial & Yeast Genomic DNA Purification kit (EURx) according to the manufacturer's instructions. Genes encoding LPMOs and BtChi_74 chitinase were PCR amplified using Pfu DNA polymerase (Thermo Fisher Scientific) and primer pairs detailed in the table below:

| Source organism | Encoded protein | Primer designation | Primer sequence (5' to 3') | Expected amplicon size (bp) | NCBI Acc. No. of the sequenced open reading frame |
|---|---|---|---|---|---|
| *Bacillus thuringiensis* | BtLPMO10_50 | BtLPMO10_50_F | AGAAGGAGATATAACT**ATG**AATAATCGATTWTTAAAACAACTACAAAAC | 1399 | OP896091.1 |
| | | BtLPMO10_50_R | GGAGATGGGAAGTCA**TTA**CACTGTTTTCCATAATGATAARGCA | | |
| | BtLPMO10_24 | BtLPMO10_24_F | AGAAGGAGATATAACT**ATG**AAAAAGAATAGTTTACARAAGRTGAAG | 697 | OP896092.1 |
| | | BtLPMO10_24_R | GGAGATGGGAAGTCA**TTA**AAATAGTGTAGGARCTTGCACTAC | | |
| | BtChi_74 | BtChi_74_F | AGAAGGAGATATAACT**ATG**AGGTCTCAAAAATTCACACTGCTAT | 2056 | OP948078.1 |
| | | BtChi_74_R | GGAGATGGGAAGTCA**TTA**GTTTTCGCTAATGACGGYATTTAAAAG | | |
| *Serratia marcescens* | SmLPMO10_21 | SmLPMO10_21_F | AGAAGGAGATATAACT**ATG**AACAAAACTTCCCGTACCCT | 625 | OP896093.1 |
| | | SmLPMO10_21_R | GGAGATGGGAAGTCA**TTA**TTTGCTCAGGTTGACGTC | | |
| | SmLPMO10_52 | SmLPMO10_52_F | AGAAGGAGATATAACT**ATG**AAATTATCCAAAATCGCCCTGAT | 1462 | OP896094.1 |
| | | SmLPMO10_52_R | GGAGATGGGAAGTCA**TTA**CTTTTTCAGGAYCCAGGCAT | | |

The amplicons were cloned into linearized pLATE11 vector, using aLICator LIC Cloning and Expression Kit 1 (Thermo Fisher Scientific) according to manufacturer's recommendations, and subsequently transformed into *E. coli* XL1 Blue. Successful constructs were sequence verified, using the Genetic Analyzer 3130x1 sequencer (Applied Biosystems) at the Laboratory of Molecular Biology Techniques at the Faculty of Biology, Adam Mickiewicz University, and the sequences of the appropriate open reading frames were deposited in GenBank, NCBI (Clark et al. 2016) and shown in the table above. The constructs harboring LPMO/chitinase-encoding fragments were subsequently transformed into *E. coli* BL21 (DE3) codon+ cells. The obtained recombinant clones were cultured in LB medium supplemented with ampicillin (final concentration: 100 µg/ml) at 37°C, until $OD_{600}$ reached 0.8-1. Next, the isopropyl β-d-1-thiogalactopyranoside (IPTG; Thermo Scientific) was added to a final concentration of 1 mM and the cultures were further incubated for 18-24 h at room temperature. The cells were pelleted in centrifuge (10 000 × g, 10 min, 4°C) and frozen in -20°C. Next, the cells were suspended in ice-cold lysis buffer (50 mM Tris/HCl, pH 8, 100 mM NaCl, 1 mM EDTA) and supplemented with lysozyme (Sigma Aldrich) and phenylmethylsulfonyl fluoride (PMSF; BioShop) to a final concentration of 0.25 mg/ml and 1 mM, respectively. One gram of cells was typically suspended in 5-8 ml of the buffer and kept 30 min on ice, with shaking (~50 rpm). Subsequently, deoxycholic acid (MP Biomedicals) was added to a final concentration of 0.8-1.3 mg/ml and ice-cold cell suspensions were sonicated with several pulses (22 kHz frequency, 14 mm amplitude), ten seconds each, using ultrasound disintegrator (UD-11; Techpan). Next, the lysates were centrifuged 40 000 × g, 10 min, 4°C and the LPMO-rich supernatant was collected. Obtained samples were then passed through 0.22 µm syringe filters (CarlRoth) and dialyzed 48 h in 14 kDa MWCO Visking tubes (Carl Roth) with 4-5 exchange of the dialysis buffer (50 mM Tris/HCl, 100 mM NaCl, pH 9). Simultaneously, as a control, the non-transformed *E. coli* BL21 (DE3) codon plus was processed using the same protocols as described above, only without antibiotic selection during culturing steps. All the obtained preparations were kept in -75°C until further use in bioassays. The protein content was verified using 10% sodium dodecyl sulfate polyacrylamide gel electrophoresis and LPMOs concentrations were estimated by densitometry with known amounts of bovine serum albumin as standards, using GelAnalyzer 2010a software (AnalystSoft Inc.). Theoretical masses of the expressed proteins were calculated upon corresponding deduced amino acid sequences using SnapGene v5.3.3 (Dotmatics).
